# Improved enzymatic labeling of fluorescent in situ hybridization probes applied to the visualization of retained introns in cells

**DOI:** 10.1101/2023.01.10.523484

**Authors:** Wen Xiao, Kyu-Hyeon Yeom, Chia-Ho Lin, Douglas L. Black

## Abstract

Fluorescence In Situ Hybridization (FISH) is a widely used tool for quantifying gene expression and determining the location of RNA molecules in cells. Here, we present an improved method for FISH probe production that yields high purity probes with a wide range of fluorophores using standard laboratory equipment at low cost. The method modifies an earlier protocol that uses terminal deoxynucleotidyl transferase to add fluorescently labeled nucleotides to synthetic deoxyoligonucleotides. In our protocol, Amino-11-ddUTP is joined to an oligonucleotide pool prior to its conjugation to a fluorescent dye, thereby generating pools of probes ready for a variety of modifications. This order of reaction steps allows for high labeling efficiencies regardless of the GC content or terminal base of the oligonucleotides. The Degree Of Labeling (DOL) for spectrally distinct fluorophores (Quasar, ATTO and Alexa dyes) was mostly greater than 90%, comparable to commercial probes. The ease and low cost of production allowed generation of probe-sets targeting a wide variety of RNA molecules. Using these probes, FISH assays in C2C12 cells showed the expected subcellular localization of mRNAs and pre-mRNAs for *Polr2a* (RNA polymerase II subunit 2a) and *Gapdh*, and of the long noncoding RNAs *Malat1* and *Neat1*. Developing FISH probe sets for several transcripts containing retained introns, we found that retained introns in the *Gabbr1* and *Noc2l* transcripts are present in subnuclear foci separate from their sites of synthesis and partially coincident with nuclear speckles. This labeling protocol should have many applications in RNA biology.

## Introduction

RNA molecules can be localized and quantified within cells using fluorescence in situ hybridization with complementary nucleic acid molecules (Chen et al. 2015; Raj et al. 2008; Femino et al. 1998). To reach single molecule sensitivity these methods employ strategies that target multiple dye molecules to the detected RNA molecule. One of the most straightforward methods is to tile several dozen complementary oligonucleotides along the target RNA, each carrying the fluorophore. These probes can be purchased from commercial vendors, but their cost limits the number of probe-sets one can test. The limited number of available fluorophores also constrains some applications.

A protocol for the enzymatic production of fluorescently-tagged DNA probes was recently described by Gaspar et al. that is inexpensive and allows efficient labeling of oligonucleotides with certain fluorophores (Gaspar et al. 2017). Some positively charged fluorophores, including ATTO488 and ATTO565, did not yield high degrees of labeling. Another limitation was that oligos with relatively high GC content yielded low labeling efficiencies presumably due to intra- and inter-molecular base-pairing interactions (Gaspar et al. 2017).

We sought to apply FISH to determine the nuclear location in mammalian cells of recently characterized classes of retained introns exhibiting different fractionation behaviors (Yeom et al. 2021). Some unspliced introns are only found associated with isolated chromatin in lysed nuclei; Other introns are present in transcripts of the soluble nucleoplasm; Still other retained introns are present in cytoplasmic mRNAs. To characterize the subcellular and subnuclear localization of these introns, we applied smFISH using enzymatically labeled oligonucleotide probe-sets. Here, we present modifications of the original protocol (Gaspar et al. 2017). With these modifications, we routinely generated probes with > 90% dye labeling. Diverse commonly used fluorophores with distinct spectra including ATTO488, ATTO565, ATTO647N, Quasar570, Quasar670; and Alexa568 could all be efficiently coupled. The coupling was insensitive to the GC content and the 3’ terminal coupled nucleotide. Using this approach, we generated dozens of probe-sets for roughly the cost of a single commercial probe-set. We then applied these probes to locate transcripts containing different classes of retained introns relative to their sites of transcription in nuclei.

## Results

### Oligonucleotides precoupled to amino-11-ddUTP are efficiently labeled with fluorophore regardless of GC content and terminal base

To address the variability in TdT labeling efficiency observed with dye-coupled ddUTP nucleotides, we changed the order of labeling steps. It was previously described that oligos carrying primary amine modifications can be labeled with succinimide-dye conjugates (Jahn et al. 2011; Winz et al. 2015; Motea and Berdis 2010), so we first added amino-11-ddUTP to the oligo 3’ ends with TdT (Fig 1A). An oligo pool targeting the exons of *Clcn6* were designed using the Stellaris probe design tool (see methods). To test the impact of GC content on ligation and labeling efficiency, we selected individual oligos from this pool ranging from 40% to 60% GC (the limits of the design tool) (Fig. S1A). We also synthesized AT and GC repeat oligos (20 mer) that were 100% AT or 100% GC. After amino-11-ddUTP addition by TdT and ethanol precipitation, the oligos were incubated with NHS-ester-Quasar570 to conjugate the dye to the terminal primary amine. These labeled oligos were again precipitated and analyzed on polyacrylamide gels (Fig. 1). As shown in Fig. 1B, the TdT modified oligos shift up in the gel, indicating the addition of one nucleotide. Across all the oligos, regardless of GC content, we observed near 100% coupling efficiency of ddUTP to the 3’ termini by the TdT enzyme (Fig.1B-1C, Fig.S1A-S1C). The conjugated dye quenched staining by SYBR-Gold causing the dye-labeled band to disappear in the gel when excited at 496 nm. These species can be observed by the dye fluorescence excited at 565 and 647 nm as a band migrating slightly above the oligo coupled to ddUTP alone. The dye labeling efficiency in the second step was consistently higher than 98% (Fig 1C) indicating the labeling is insensitive to GC content.

**Figure 1.**
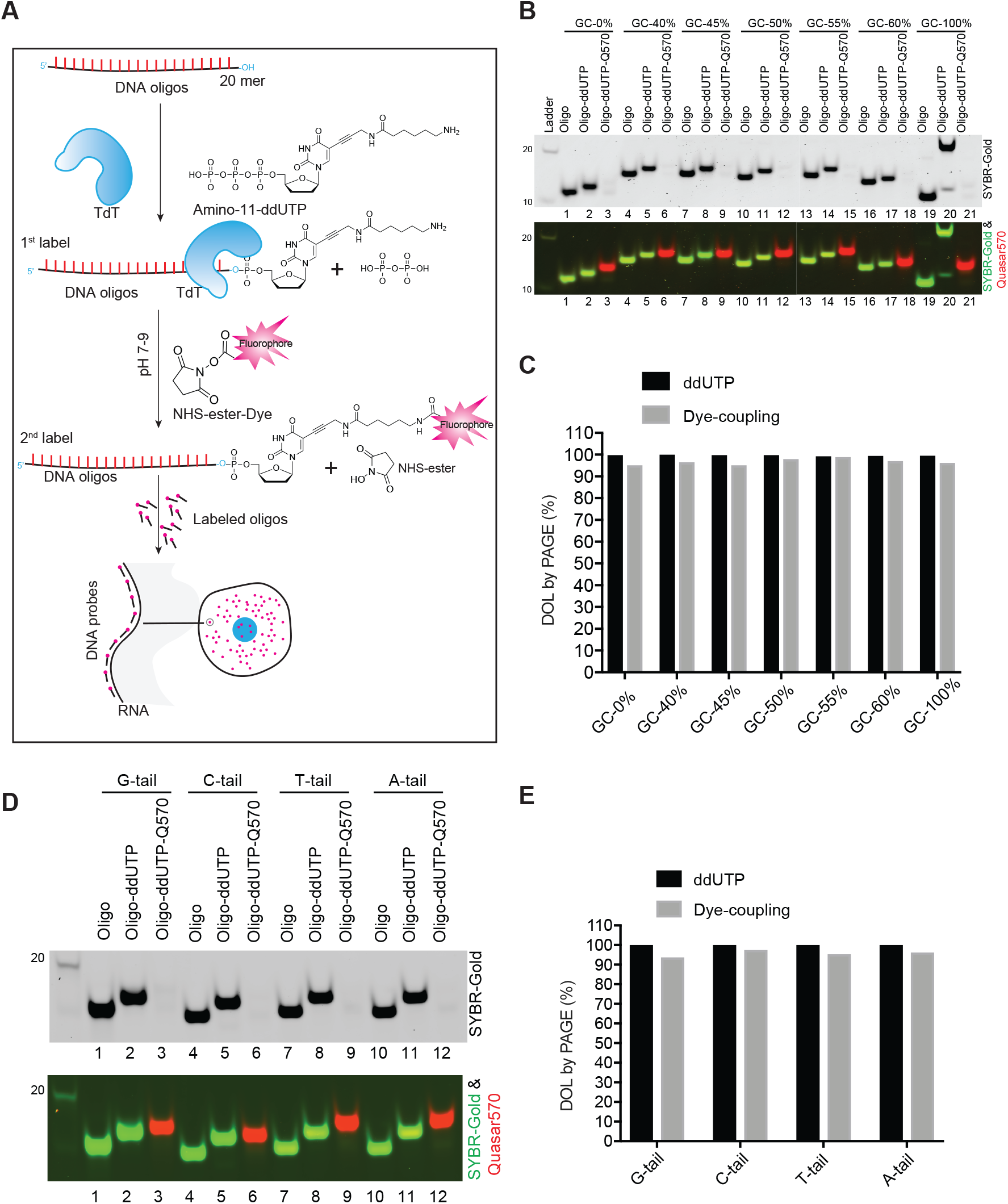
Efficient oligo coupling of amino-11-ddUTP and post-labeling with NHS-ester dyes. **(A)** Diagram of the probe labeling workflow. A set of single stranded oligonucleotides (primers) are first coupled to amino-11-ddUTP at their 3’ end using the TdT enzyme. After ethanol precipitation, amino-11-ddUTP coupled oligos are labeled with an NHS-ester conjugated fluorescent dye. After ethanol precipitation and column purification, the oligos are used for hybridization to cells and microscopic imaging. **(B)** PAGE analysis of the Degree Of Labeling (DOL) of ddUTP and dye for oligos of different GC content. Upper panel: SYBR-Gold channel in grey. Bottom panel: the composite of the SYBR-channel (green) and the Quasar 570 channel (red). **(C)** Quantification of Fig. 1B. **(D)** PAGE analysis of 1^st^ and 2^nd^ step labeling efficiency for oligos with 4 different 3’ ends. Upper panel: SYBR-Gold channel in grey. The bottom panel shows a composite of the SYBR-channel (green) and the Quasar 570 channel (red). (**E)** Quantification of Fig. 1D.

TdT is known to be relatively insensitive to the 3’ terminal nucleotide it is modifying (Winz et al. 2015; Motea and Berdis 2010). To assess whether variation of the last base influenced the dye labeling, we generated the 50% GC content oligo described above with each of the four nucleotides (A,G,C,T) appended to the 3’ end. We found that the last base had little effect on either the ddUTP addition by TdT or the dye conjugation, with all four variants yielding a degree of labeling above 90% (Fig. 1D-1E and Fig. S1D-S1F). These experiments were repeated using Quasar670 in place of Quasar570 with equivalent results, showing that two standard dyes can be used in this procedure (Fig.1, Fig. S1). These results indicate that by switching the order of addition for the ddUTP and the dye one can obtain a high degree of probe labeling regardless of GC content and terminal base.

### Pools of oligonucleotides can be labeled with equivalent efficiency to commercial FISH probe sets

A typical FISH probe set contains several dozen oligonucleotides and it is impractical to label these individually (Young et al. 2020; Raj et al. 2006). We next examined the labeling efficiency for oligo pools designed as FISH probe sets to a variety of targets. These probe sets contained 24 to 48 different oligos mixed in equimolar concentrations. As mixtures of sequences these probe sets migrate as more diffuse bands than single oligos (Figure 2). The oligo mixtures were then modified with Amino-11-ddUTP using TdT and conjugated to NHS-Quasar570 or NHS-Quasar670 as above. We labeled 14 different oligo sets with Quasar570 (Sets A to N, Fig. 2A, Table S3). We labeled 10 oligo sets with Quasar670 (Sets I to VI, VI to XI, Fig. 2D-2F, Table S4). For comparison we purchased two Stellaris probe sets targetting the exons of Gapdh and the noncoding RNA-Lexis (Fig. 2A, 2D). We used two methods to quantify the Degree of Labeling (DOL). We measured the absorbance of the dye and oligo by spectrophotometry and calculated their relative concentrations using the extinction coefficient for dye and the estimated extinction coefficient for the oligo, as done previously (Gaspar et al. 2017). As an alternative approach, we also quantified the intensities of the bands on PAGE gels and determined the DOL by comparing the SYBR gold intensity of remaining unlabeled probes after labeling to the intensity of original oligos before labeling (Fig. S2). Across the 25 probe sets and for both dyes, these two methods yielded very similar results, with probe sets exhibiting an average DOL values of 95% and 98% in PAGE and spectrophotometric analysis respectively (Fig. 2A-2B, 2D-2E). By PAGE, we measured DOL values of 88% and 94.5% for the Stellaris Lexis and Gapdh probes, respectively, indicating that our labeling protocol is at least comparable (Fig. 2A, 2C, 2D and 2F).

**Figure 2.**
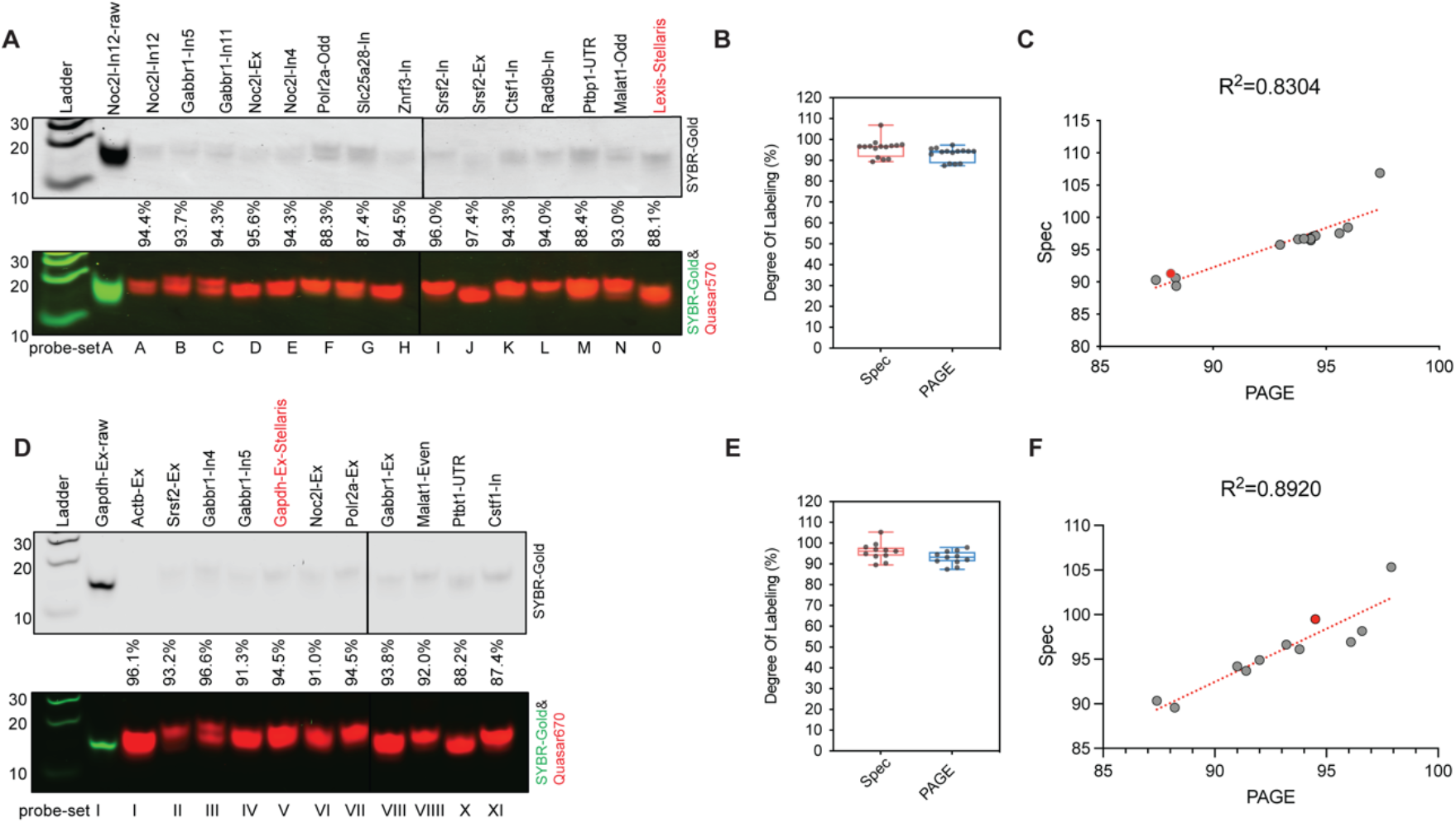
Complete FISH probe sets can be efficiently labeled in bulk with comparable efficiency to commercial probes. **(A)** PAGE analysis of Quasar570 labeling of multiple probe sets. Probe-sets A to O (lanes 3-17) contained from 15 to 48 oligonucleotides each, targeting a particular cellular transcript. Lane 2 contains the unlabeled oligos of set A. Lane 17 (set O) contains a commercial probe set from Stellaris. Band Doublets in some lanes indicate mobility differences among the mixed probes. **(B)** Box plots show the DOL of multiple probe sets determined by spectrophotometer and by PAGE analysis. Each dot represents one probe set. **(C)** The correlation of DOL measurements for Quasar 570 labeling by spectrophotometer and PAGE analysis. Probe sets from Stellaris are highlighted in red. **(D)** PAGE analysis of Quasar670 labeling efficiency for multiple oligo sets. Sets I to XI (lanes 3 to 13) are dye-labeled oligo sets containing 15 to 48 oligos each. Lane 2 contains the original unlabeled oligos of set I. Lane 7 (set V) contains a commercial probe set from Stellaris. **(E)** Box plots of the Quasar 670 DOL for the multiple probe sets determined by spectrophotometer (see Fig. S2) and PAGE. **(F)** The correlation of Quasar 670 DOL measurements by spectroscopy and PAGE is displayed by scatter plot, with R square value indicated.

### Labeling is compatible with a range of dyes

Visualization of multiple RNA species within a single cell requires the probe-set for each target to carry a dye with a different emission wavelength (Young et al. 2020). Ideally one would have a stock of Amino-11-modified oligos that could be coupled to different dyes as needed. We next tested a variety of spectrally distinct NHS-modified fluorophores for their efficiency of coupling to the probe-sets described above for the Quasar dyes. These fluorophores included ATTO-488, ATTO-565, ATTO647N, Alexa-568 and Alexa-647. The average DOL for all the ATTO dyes was >95% according to both PAGE and spectrophotometer analysis (Fig. 3A–3B, Fig. S3A-S3C). Alexa568 had an average DOL >90% similar to the ATTO dyes (Fig. 3A–3B, S3D). In contrast, Alexa647 dye performed poorly with an average DOL of ~20% in these labeling conditions (Fig. 3A, 3B, S3E). Doubling the concentration of NHS-ester-Alexa647 in the labeling reaction increased the DOL to 35%. This NHS-ester-dye conjugate appears to be inherently less reactive than the others (Fig. 3A–3B, S3E). To test the correlation of the two analytical methods, we plotted and fitted the spectroscopic and densitometric DOL to a linear regression model (Fig. S4A-S4F). As seen for the Quasar dyes, quantifying the DOL of the ATTO and Alexa dyes by either PAGE or spectrophotometry gave similar results. Thus, the labeling protocol functions well with two Quasar dyes, three ATTO dyes and one Alexa dye, allowing use in multi-channel FISH assays.

**Figure 3.**
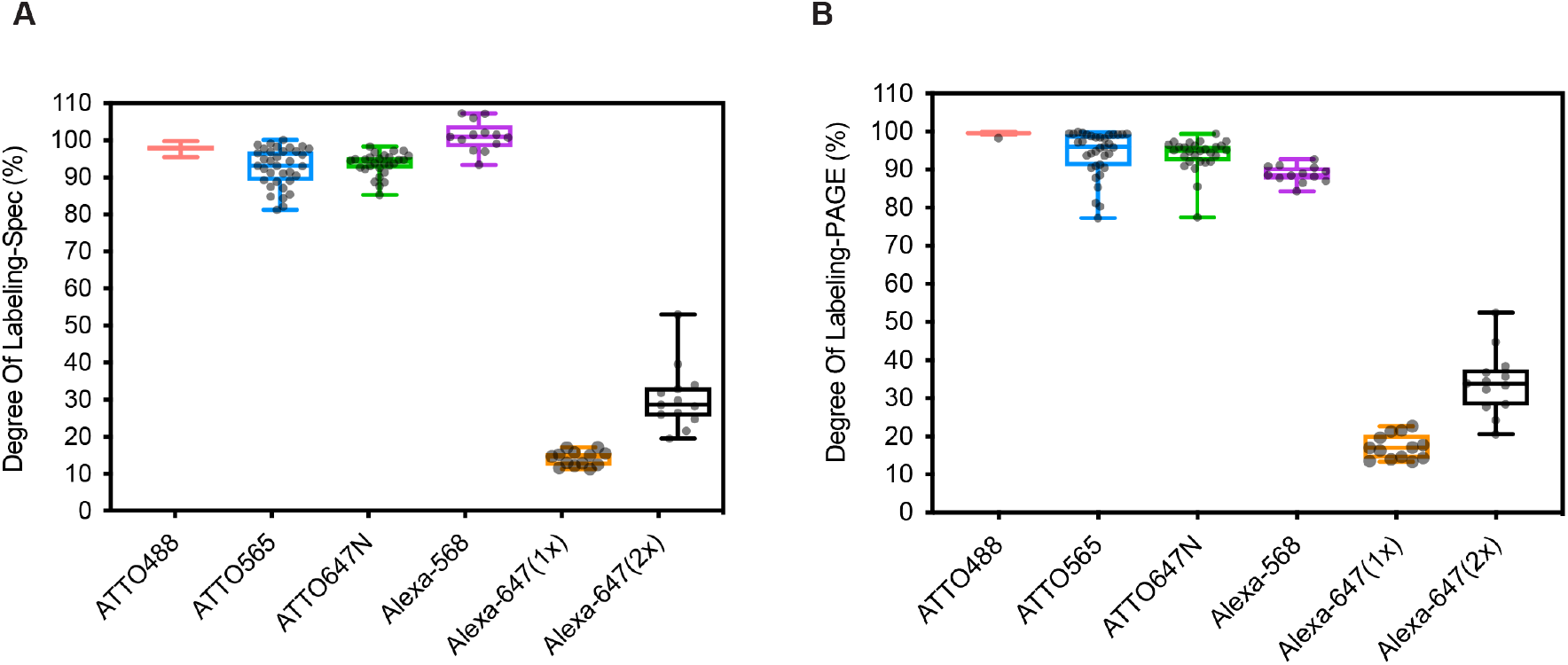
Labeling efficiency for spectrally distinct dyes. **(A)** Box plots show the degree of labeling of multiple probe sets using a series of ATTO and Alexa dyes determined by spectrophotometer. **(B)** Box plots of the DOL for ATTO and Alexa dye labeling determined by PAGE. Each dot represents one probe set containing up to 48 oligos (All sequences are in Tables S5-S9).

### Performance of the probes in FISH assays

To test our probes in smFISH, we designed probe sets against a variety of cellular RNAs with previously published patterns of localization, including Polr2a, Malat1, Neat1, and Gapdh. These were designed using the Stellaris tool, synthesized and labeled as above. The Polr2a transcript (RNA polymerase II subunit A) was chosen for its moderate expression and exon length (6.7kb) that can accommodate a large number of probes (Xie et al. 2018). The 48 oligonucleotides of the Polr2a exon probe-set were split into odd and even subsets (24 oligos each) that were coupled to different Quasar dyes (Quasar570 and Quasar670 respectively; Fig. 4A). The odd and even probes with different fluorophores were combined prior to FISH hybridization. FISH images acquired by confocal microscopy showed odd and even spots predominantly dispersed in the cytoplasm of the C2C12 cells (Fig. 4A). Combining the odd and even channels showed visually that these spots almost entirely colocalized (Fig. 4A). Quantifying the intensities for each pixel in ImageJ and comparing them using the JACoP plugin generated a Pearson Correlation coefficient of 0.774 (Fig. S5). The numbers of foci observed with the odd and even probes were also quantified using the program FISH-quant (Mueller et al. 2013). The per cell spot numbers for the odd and even probes of Polr2a were very similar (odd: Mean=103 and even: Mean=97) (Fig. 4B), and highly correlated across many cells (R^2^= 0.955; Fig. 4C). This pattern of FISH staining is nearly identical to that published previously and indicates that the probes function well to quantify mRNA within cells (Xie et al. 2018).

**Figure 4.**
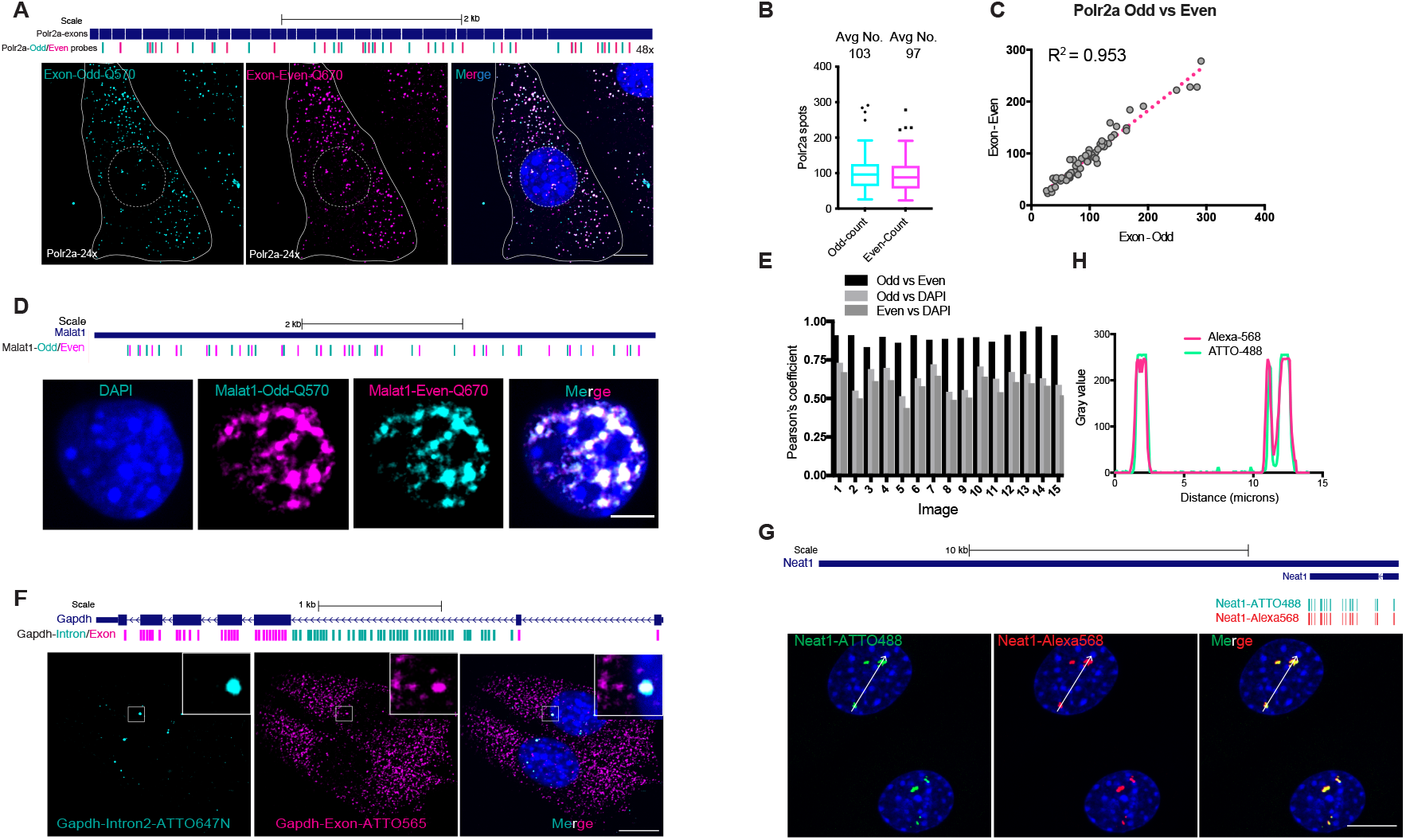
The performance of fluorescently labeled probes in FISH assays. **(A)** Above: diagram showing the locations of Odd and Even probes across the Polr2a coding regions (exons) in the mouse genome; Below: Confocal microscopy images of C2C12 cells probed with the Polr2a Odd and Even FISH probes. The nuclear outline is indicated by a dashed line, and the cell boundary by a solid white line. **(B)** Box plots of the spot counts in C2C12 cells for Polr2a in the Odd and Even channels using FISH-quant software. The average number of spots per cell for the Polr2a Odd and Even signals is shown above. **(C)** The correlation of the odd and even spot numbers per cell are displayed in a scatter plot with the R^2^ indicated. 59 cells were counted and plotted. **(D)** Above: Diagram of the locations of the Odd and Even probes for Malat1. Below: Confocal Microscopy images of C2C12 cells probed with the Malat1 Odd and Even probes. **(E)** Bar graph of Pearson’s coefficients determined for each of 15 images. Pairwise correlations of the per pixel fluorescence intensity in two channels: Quasar570 (Malat1-Odd) vs Quasar670 (Malat1-Even), Quasar570 (Malat1-Odd) vs DAPI, and Quasar670 (Malat1-Even) vs DAPI. **(F)** Above: Diagram of the locations of Exon (magenta) and Intron (cyan) probes for Gapdh arrayed along the annotated mouse gene. Below: Confocal microscopy images of C2C12 cells hybridized with the Gapdh exon and intron probes. The white box highlighted area are zoomed in on the upright corner accordingly. **(G)** Above: Diagrams of the ATTO488 (green) and Alexa568 (red) probe locations arrayed along the Neat1 short isoform in the mouse gene annotation. Below: Confocal microscopy images of C2C12 cells hybridized with 25 Neat1 ATTO488 probes (Green), and 25 Neat1 Alexa568 probes (Red), with DAPI in blue. **(H)** Line scan profile of the ATTO488 (green) and Alexa568 (red) channels for the cell shown in G. Scale bar = 10 microns.

The Quasar dyes also performed well for localizing the long noncoding RNA Malat1. We created Malat1 odd (Quasar570) and even (Quasar670) probes (Fig. 4D). Malat1 is abundantly expressed and too concentrated in nuclear speckles discern single molecule spots for counting in FISH-quant (Fig. 4D)(Fei et al. 2017; Wang et al. 2021). We measured fluorescence intensities for the odd and even channels for each pixel across the area of the nucleus in ImageJ. These per pixel intensities for the odd and even probes were highly correlated, with an average Pearson’s Coefficient of 89.4%. As expected, the Malat1 probe signals in speckles were less correlated with DAPI, with the Odd vs DAPI comparison yielding a Pearson’s Coefficient of 63%, and the Even vs DAPI yielding 56.6% (Fig. 4E).

We next tested the use of dual probes to different portions of the same transcript. Gapdh RNA has been shown in previous studies to exhibit abundant mRNA in the cytoplasm (Rowland et al. 2019). In contrast, Gapdh introns are excised rapidly at the site of transcription in the nucleus (Rowland et al. 2019; Femino et al. 1998). Using ATTO565 labeled probes to target the exons and ATTO647N probes to label a long Gapdh intron, we confirmed this pattern (Fig. 4F). The intron probes produce two small foci per nucleus corresponding to the transcribed genes. The exon probes label these nuclear spots as well as abundant cytoplasmic mRNA (Fig. 4F). To test the Alexa568 dye, we generated two sets of probes to the lncRNA Neat1. Neat1 is a component of nuclear paraspeckles and exhibits a distinctive staining pattern with several prominent nuclear foci (Ahmed et al. 2018). Probing with Neat1 oligos labeled with both ATTO488 and Alexa568 produced the expected 2 to 4 foci per nucleus (Fig. 4G). The signals for ATTO488 and Alexa568 almost entirely overlapped, with the two dyes performing equivalently (Fig. 4H).

These experiments indicate that probes generated by this method perform well in FISH experiments using tissue culture cells.

### Transcripts containing retained introns form nuclear foci distant from the site of transcription

Our goal in optimizing the FISH method was to determine the nuclear location of transcripts containing retained introns. In earlier studies, we defined different classes of retained introns based on their fractionation behavior from lysates of mouse embryonic stem cells (ESC), neuronal progenitor cells, and cortical neurons. Several classes of introns found in nuclear polyadenylated RNA of ESC were distinguished from classical retained introns found in cytoplasmic mRNAs (Yeom et al. 2021). Some transcripts, such as Noc2l pre-mRNA, contain unspliced introns in the chromatin fraction and/or the soluble nucleoplasm but are expressed as fully spliced mRNAs in the cytoplasm (Fig. 5A). These can be classified as “detained” introns in that they are abundant in the nucleus, but are ultimately excised and the RNAs in the cytoplasm are fully spliced (Shah et al. 2018a; Yeom et al. 2021; Boutz et al. 2015; Braun et al. 2017; Pandya-Jones et al. 2013; Mauger et al. 2016). In contrast, other polyadenylated pre-mRNAs in ESC, such as Gabbr1, exhibit different fractionation behavior that is similar to unpolyadenylated nascent transcripts and certain long noncoding RNAs. These tanscripts are tightly associated with the pelleted chromatin, but found only in limited amounts in the soluble nucleoplasm and cytoplasm (Fig. 5E)(Yeom et al. 2021). Introns in these RNAs remain unexcised and the transcripts do not produce mature mRNA. Speculating that these RNAs might be anchored at their sites of transcription, we applied FISH to examine to the location of nuclear polyadenylated RNAs containing highly retained introns.

**Figure 5.**
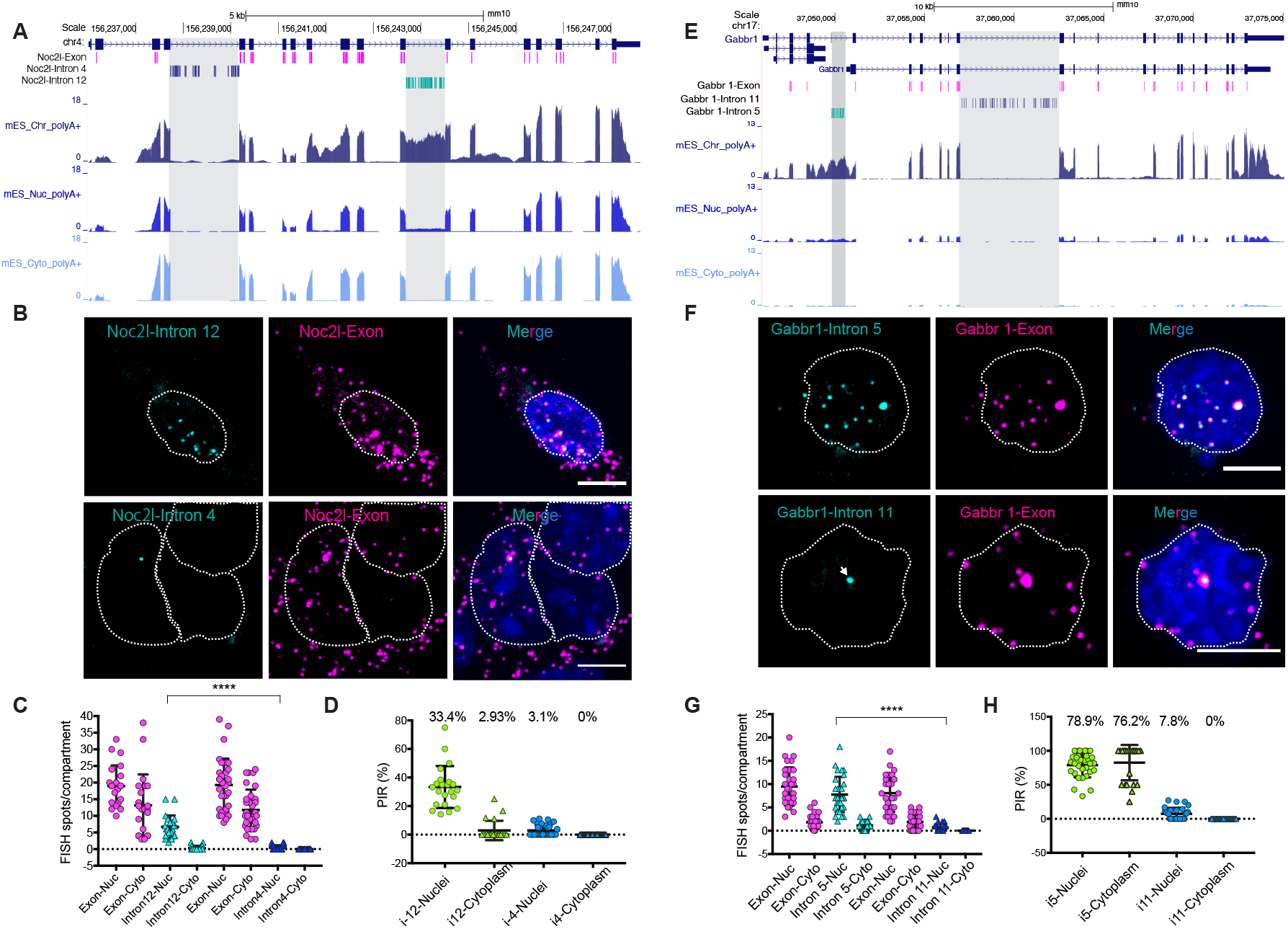
Localization of retained introns in mouse ES cell nuclei. **(A)** UCSC genome browser tracks show Noc2l polyA^+^ RNA from the chromatin, nucleoplasmic, and cytoplasmic fractions. Top: the genomic structure of the Noc2l gene is diagrammed with exon and intron probe locations in magenta, blue and green. Bottom: RNA seq reads from the three fractions are displayed. Introns probed by FISH are indicated in gray. **(B)** FISH images of mES cells probed for Noc2l introns 12 (ATTO647N, retained), 4 (ATTO647N, unretained) and exons (ATTO565). Nuclear borders are outlined in white dashed lines. **(C)** Quantification of spot counts for the Noc2l exon and intron probes in the nuclear and cytoplasmic compartments. **(D)** Percentage intron retention (PIR) for Noc2l introns 12 and 4 in nuclear and cytoplasmic compartments as determined by the ratio of intron to exon spot numbers in mES cells. **(E)** UCSC genome browser tracks show Gabbr1 polyA^+^ RNA from the chromatin, nucleoplasmic, and cytoplasmic fractions. Top: the genomic structure of the Gabbr1 gene is diagrammed with exon and intron probe locations in green, blue and magenta. Bottom: RNA seq reads from the three fractions are displayed. Introns probed by FISH are indicated in gray. **(F)** FISH images of mES cells probed for Gabbr1 introns 5 (ATTO647N, retained) and 11 (ATTO647N, unretained) in green and for the Gabbr1 exons (ATTO565) in magenta. Nuclear borders are outlined in white dashed lines. **(G)** Quantification of spot counts for the Gabbr1 exon and intron probes in the nuclear and cytoplasmic compartments. **(H)** Percentage intron retention (PIR) for Gabbr1 introns 5 and 11 in nuclear and cytoplasmic compartments as determined by the ratio of intron to exon spot numbers in mES cells. Scale bar = 5 microns.

To examine spliced and unspliced Noc2l transcripts in ESC, we generated separate FISH probe-sets for the Noc2l exons, and for introns 4 and 12 (Fig. 5A, 5B). Intron 4 is highly spliced in all compartments, whereas intron 12 is largely unspliced in the chromatin fraction, present at lower levels in the soluble nucleoplasm, and fully excised in the cytoplasm. Mouse ES cells probed for the Noc2l exons showed abundant spots in the cytoplasm as expected, as well as multiple foci in the nucleus. Cells probed for the rapidly spliced intron 4 showed only 0, 1 or 2 nuclear foci that also stained with exon probes, and are presumably the sites of transcription (Fig. 5A-5B, Fig. S6B). Intron 4 staining was not observed in the cytoplasm in agreement with the sequencing data (Fig.5A, Fig.S6B). In contrast to intron 4, the probes for the highly retained intron 12 colocalized with nearly all the nuclear exon foci whether they were at the sites of transcription or not. These intron 12 foci were absent from the cytoplasm (Fig. 5C, 5D, Fig. S6A). Thus, transcripts containing this retained intron do not remain at the site of transcription but instead are spliced after release from the gene locus but before export to the cytoplasm. This is in agreement with prior studies of introns that are spliced post-transcriptionally (18–21).

We next examined the localization of the Gabbr1 transcripts that exhibit a different pattern of splicing and fractionation. In mouse ESC, Gabbr1 transcripts are almost entirely unspliced in a region of complex processing encompassing intron 5 (Fig. 5E). Other Gabbr1 introns including intron 11 appear spliced even in the chromatin fraction. We generated Gabbr1 probes for the exons, for intron 11, and for the region of intron 5 present in the major isoform expressed in mESC (Fig. 5E). All three probe sets yielded very little staining in the cytoplasm in agreement with the fractionated RNAseq data indicating that these RNAs remain in the nucleus. The limited spots observed in the cytoplasm usually stained for both the exons and intron 5. Exon foci for Gabbr1 were more abundant in the nucleus, with most cells exhibiting one large spot per nucleus and 5-15 smaller spots. Nearly all these exon foci costained for intron 5 (Fig. 5F, Fig. S6C). In contrast, the probes for the rapidly excised intron 11 yielded only one spot per nucleus, which is presumably the site of transcription (Fig. 5F, Fig. S6D). This was often the largest exon spot in the nucleus and costained with the exon and intron 5 probes. The spot numbers for introns 5 and 11 of Gabbr1 and for introns 4 and 12 of Noc2l in nucleus were quantified (Fig. 5C, 5G). The rapidly excised introns 12 and 4 averaged less than 2 foci per nucleus, consistent with them marking the gene loci. In contrast, Noc2l intron 11 and Gabbr1 intron 5, which exhibit much lower levels of excision, generated many more nuclear foci, whose numbers approached the numbers of exon foci. Computing the Percent Intron Retention (PIR) from the ratio of intron to exon foci yielded values in agreement with those calculated from sequencing data (Fig. 5D, 5H, Fig. S6E). Although the foci at the transcriptional loci were larger, Gabbr1 RNA localization was similar to Noc2l, with multiple foci scattered through the nucleus away from the site of transcription (Fig. 5G–5H). These data indicate that despite differences in their fractionation behavior and eventual splicing, RNAs containing retained introns are being released from both of these gene loci. In the case of Noc2l, these RNAs are eventually spliced and exported to the cytoplasm, whereas for Gabbr1 the RNAs remain unspliced and unexported. The Gabbr1 RNAs are also more strongly associated with the chromatin pellet fraction after solubilization in stringent buffers (Yeom et al. 2021).

### Highly retained introns partially coincident with nuclear speckles

Nuclear speckles are regions within the nucleoplasm enriched for splicing factors and for the long noncoding RNA Malat1. Active gene loci are often observed at the periphery of speckles and posttranscriptional splicing is observed within them (Girard et al. 2012; Dias et al. 2010; Galganski et al. 2017). Using Malat1 as a speckle marker, we examined whether the retained intron transcripts are within speckles. For Noc2l and Gabbr1, we used separate labels for the retained intron and exon probes with a third fluorophore on the Malat1 probes in multiplexed FISH assays. Identifying foci of unspliced introns defined by colocalized exon and intron sequences, observing ~5 colocalized exon-intron spots for Noc2l per nucleus and ~10 such spots for Gabbr1 (Fig. 6A–6B, 6D–6E). We then measured the overlap of these retained intron foci with Malat1 staining in speckles. We found that the majority of these retained intron transcripts colocalized with Malat1 (Fig. 6A–6B, 6D–6E). On average, 70% of the Noc2l-intron 12 spots and 76% of the Gabbr1-intron 5 overlapped with Malat1 staining (Fig. 6C and 6F). However, for both introns there were a subset of transcripts that did not appear to be in speckles. Simultaneous probing for Noc2l-intron12 and Gabbr1-intron5 did not show overlap (Fig. 6G), indicating that speckles are not homogeneous but instead contain different RNAs in different regions.

**Figure 6.**
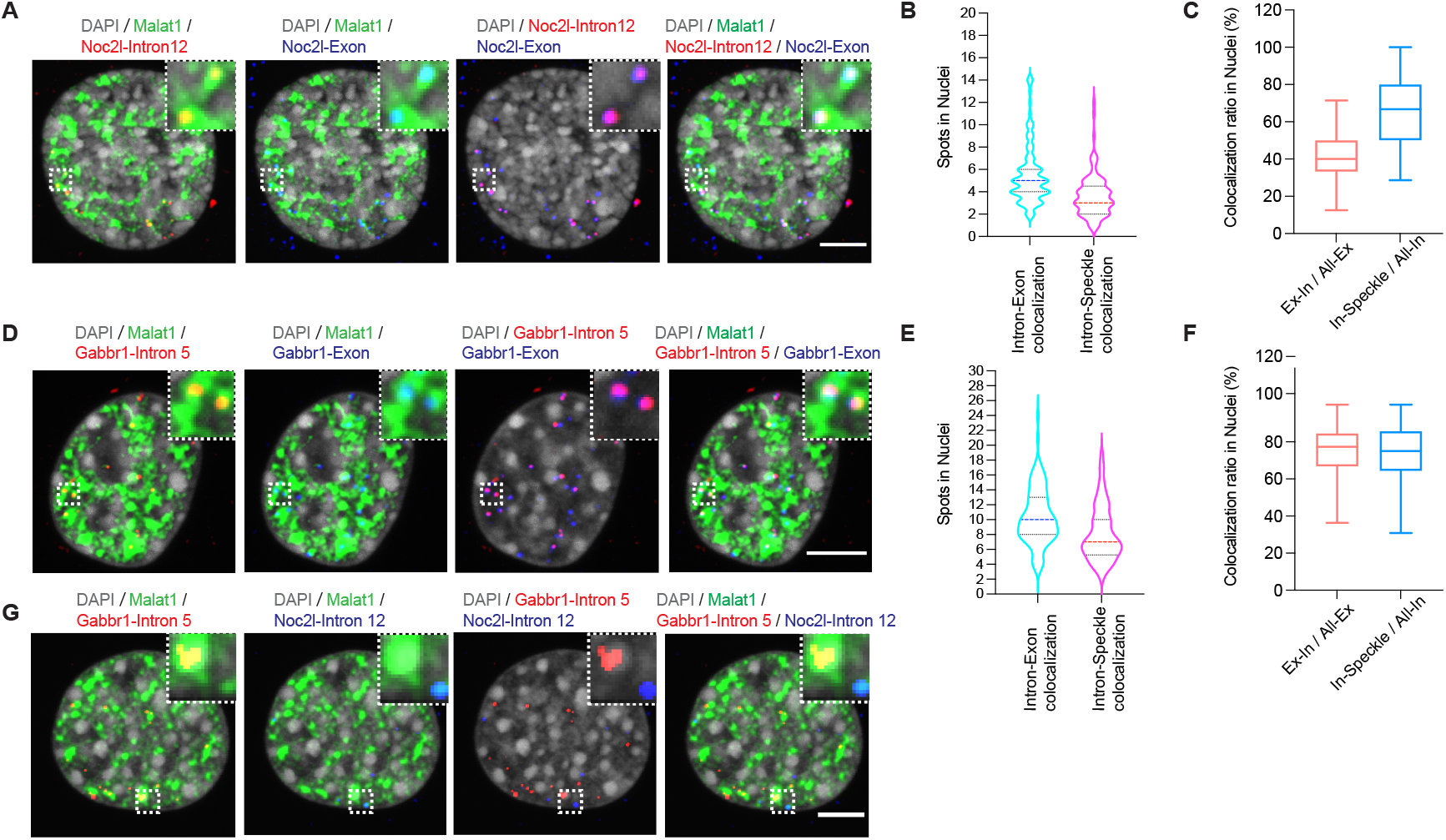
Retained intron transcripts of Gabbr1 and Noc2l are dispersed in the nucleus and partially colocalize with speckles. **(A)**. FISH images of C2C12 cells simultaneously probed for Malat1 (ATTO488, green), Noc2l-intron12 (ATTO647N, red), and Noc2l-exon (ATTO565, blue). DAPI stain is shown in grey. **(B)** Violin plots showing numbers of colocalized Noc2l exon and intron spots, and numbers of intron spots colocalized with speckle signal, across 101 cells. **(C)** Box plots showing the ratios of the colocalized Noc2l intron-exon spots over the total exon spots in nucleus(In-Ex / All-Ex), and the intron spots with overlapping speckle (Malat1) signal over the total intron spots in nucleus(In-Speckle / All-In), across 101 cells. **(D)**. FISH images of C2C12 cells simultaneously probed for Malat1 (ATTO488, green), Gabbr1 intron 5 (ATTO647N, red), and Gabbr1 exons (ATTO565, blue). DAPI stain is shown in grey. **(E)** Violin plots showing numbers of colocalized Gabbr1 exon and intron spots, and numbers of intron spots colocalized with speckle signal, across 104 cells. **(F)** Box plots showing the ratio of the colocalized Gabbr1 intron-exon spots over the total exon spots in nucleus, and the intron spots with overlapping speckle (Malat1) signal over the total intron spots in nucleus, across 104 cells. **(G)** FISH images of C2C12 cells simultaneously probed for Malat1 (ATTO488, green), Gabbr1-intron5 (ATTO565, red), and Noc2l-intron12 (ATTO647N, blue), with DAPI stain in grey.

## Discussion

### Labeling oligonucleotide probe sets

We present an improved method for the rapid, efficient and inexpensive production of fluorescent oligonucleotides. This protocol is derived from an earlier one using terminal deoxynucleotidyl transferase to add functional moieties to the 3’ end of a standard DNA oligo (Gaspar et al. 2017). We found that switching the order of addition and making several smaller adjustments, significantly improved the labeling efficiency and allows the use of a broader range of fluorophores and oligonucleotide sequences.

In the new procedure Amino-11-ddUTP is first coupled to an oligonucleotide pool using TdT. This step is very efficient and allows the creation of a stock probe-set that can be subsequently labeled with the fluorophore of choice. The oligo pools are then treated with a succinimide-dye conjugate to add the fluorophore. For nearly all dyes tested, this labeling is also very efficient, and not affected by either the GC content or 3’ terminal nucleotide of the probe. Comparing Quasar dye-coupled lab-made probes with commercial Quasar probes, we found that they had comparable labeling efficiencies and performed equivalently. Besides the Quasar dyes, we successfully labeled probes with commonly used ATTO dyes (ATTO488, ATTO565 and ATTO647N) and an Alexa dye (Alexa568) enabling many combinations for multiplexed imaging (Fig. S3-S4, Table S5-S9). One dye, Alexa647, did not efficiently couple with the amino group and was not usable. It will be interesting to test additional dyes and other modifications such as biotin for coupling to our oligonucleotide probe sets.

In the course of these experiments, we generated more than 100 probe sets containing 24 to 48 oligos each and labeled with six different dyes. With these materials the labeling of one probe-set costs about 15 dollars, and can be used in approximately 1000 FISH assays (Table S1). This is compared to the approximate $800 cost of a single commercial probe set of 48 oligonucleotides labeled with a Quasar dye.

The use of tiled oligonucleotides to target multiple fluorophores to an RNA molecule by in situ hybridization has been a standard strategy for in vivo imaging experiments. Several other cost-effective FISH probe protocols have been developed. The single molecule inexpensive FISH (smiFISH) uses unlabeled primary probes against the RNA target along with dual fluorescent probes binding to the primary probe for signal detection (Tsanov et al. 2016; Calvo et al. 2021). Another FISH method, termed single molecule hybridization chain reaction (smHCR), employs unlabeled primary probes to the target RNA and then chain reaction probes that can be labeled and amplified (Shah et al. 2016). Another enzyme-based FISH method employs phagemid to produce ssDNA probes. The purified and DNase I digested ssDNAs (normally 20-60 nt) are then used for enzyme-based probe synthesis (Lanctôt 2018). These methods each have their own potential advantages. It will be useful to compare them to standard oligonucleotide FISH now that their costs are more equivalent.

### Localizing retained introns

Most introns are excised during transcription of the nascent RNA and localize to the gene locus in FISH analyses (Shah et al. 2018b). Other introns are excised after transcription is complete. Some of these unspliced transcripts are released from the DNA template and can be observed in the nucleoplasm at a distance from the gene and sometimes in the cytoplasm (Coté et al. 2020; Girard et al. 2012; Vargas et al. 2011; Coulon et al. 2014). In earlier studies, we and others have characterized retained introns and their regulation (Monteuuis et al. 2019). Classical retained introns are found in transcripts that have been exported to the cytoplasm as mature mRNAs. Other introns exhibit delayed splicing and are abundant in nuclear RNA but are excised prior to export of the fully processed RNA. These “detained” introns have been observed in inflammatory response transcripts of mouse macrophages, in transcripts affecting cell cycle regulation and growth control in human cell lines, in cultured postmitotic neurons and in mouse brain (Bhatt et al. 2012; Boutz et al. 2015; Mauger et al. 2016; Pandya-Jones et al. 2013; Frankiw et al. 2019; Wong et al. 2013). These introns are thought to act as brakes that slow the expression of the transcripts where they are found, and in some cases to create a pool of RNAs that can be rapidly matured into mRNA in response to a signal. In different systems, reduced excision rates have been ascribed to reduced activity of the Arginine Methyltransferase PRMT5, and to the presence or absence of particular RNA binding proteins (Braun et al. 2017; Hayashi et al. 2014; Frankiw et al. 2019). Other studies found that one class of retained introns was derived from genes associated with the nuclear periphery and lamina, whereas another class including most detained introns was expressed from more centrally located genes whose transcripts colocalized to nuclear speckles (Barutcu et al. 2022; Tammer et al. 2022).

We recently characterized retained introns based their fractionation into cytoplasmic and two subnuclear compartments (Yeom et al. 2021). Within the nuclei, we identified both nascent and polyadenylated RNAs that were tightly associated with the high molecular weight pellet fraction containing chromatin and some of the nuclear speckle material. This fraction was distinguished from a soluble nuclear fraction that contains mature mRNAs in transit to the cytoplasm, as well as some polyadenylated but incompletely spliced RNAs. Nuclear introns included the described detained introns that were found both in the chromatin pellet and the soluble nucleoplasm, but were absent from mature RNA in the cytoplasm.

We also identified an unusual class of nuclear polyadenylated transcripts that contain introns but are absent from the cytoplasm as either spliced or unspliced RNA. These transcripts were tightly associated with the chromatin pellet, showing fractionation behavior similar to Xist and other chromatin bound lncRNAs (McHugh et al. 2015; Pandya-Jones et al. 2020; Mishra and Kanduri 2019; Yeom et al. 2021). For these RNAs, the failure to splice and possibly their tight association with chromatin prevents expression of the encoded protein. The most notable example is Gabbr1, which is transcribed in ESC but fails to splice intron 5; Gabbr1 transcripts also remain tightly associated with the chromatin pellet and, despite substantial transcription, fail to produce Gabbr1 protein. As cells differentiate into neurons, these Gabbr1 gene RNAs become fully processed to produce cytoplasmic mRNA and Gabbr1 protein. The unusual fractionation behavior of the Gabbr1 transcripts in ESC and its similarly to Xist, led to the hypothesis that these transcripts might condense at their site of transcription rather than being released into the nuclear speckle as seen with RNAs destined to be fully spliced, such as detained intron transcripts.

To examine the location of Gabbr1 transcripts we developed FISH probe sets for the retained Gabbr1 intron 5, for the Gabbr1 exons, and for Gabbr1 intron 11. Intron 11 is excised cotranscriptionally and unlike intron 5 is only seen in the nascent unpolyadenylated Gabbr1 RNA. The localization patterns of the Gabbr1 RNAs were compared to those of Noc2l. Noc2l intron 12 is also highly retained in the nuclear RNA but is eventually excised and, unlike Gabbr1, the mature Noc2l mRNA is exported and translated. The localization of Noc2l intron 12 was compared to the Noc2l exons and Noc2l intron 4, which is cotranscriptionally excised. Probing for the cotranscriptionally excised introns identified the sites of the active gene loci, which also labeled with exon and retained intron probes for both genes. In contrast to the cotranscriptional introns, the retained introns for both the Gabbr1 and Noc2l genes, were also seen in foci distant from the sites of transcription that co-labeled with the exon probes. Thus, both of these retained intron transcripts are being released from the DNA template. The Noc2l transcripts will eventually be spliced but the Gabbr1 transcripts are presumably destined for degradation. The signal for these transcripts partially colocalizes with the speckle marker Malat1, consistent with previous results showing retained introns in speckles. Interestingly, the foci at the transcription site for Gabbr1 are noticeably larger than seen for Noc2l or other transcripts, possibly an indication that it forms a different molecular condensate.

## Materials and Methods

### FISH probe production and purification

#### 1. Oligo design and purchase

Arrays of 20-mer DNA oligos complementary to RNA targets were designed using the Stellaris Probe Designer version 4.2 (https://www.biosearchtech.com/stellaris-designer). Selection criteria included 40%-60% GC content and similar melting temperatures. The spacing length was at least 2 nucleotides between oligos. Oligos were purchased from IDT as desalted oligos in 96 well plates at 200 μM concentration in nuclease-free water. Oligos were thawed and spun to remove debris before labeling.

#### 2. Labeling oligo pools with Amino-11-ddUTP

Probe set pools were assembled by mixing 10 μL from each oligo well. Enzymatic labeling was performed in a 40 μL mixture containing: 28 μL of the oligo pool (200 μM in H_2_O), 1 μL Amino-11-ddUTP (10 mM in H_2_O, Lumiprobe), 8 μL of 5x TdT buffer (Thermo scientific), 3 μL TdT enzyme (20 U/μL, Thermo scientific). Labeling reactions were incubated at 37C overnight in a PCR thermocycler with a hot lid. Reactions were stopped by incubation at 70c for 10 min. The oligos were then ethanol precipitated by addition of 4 μL sodium acetate (3M, pH5.2, AMRESCO, Cat.E521-100 ml) and 110 μL 100% ethanol and cooled to −80c for at least 1 h. The samples were then centrifuged at 14,000 rpm for 15 min to pellet the oligos. The oligo pellets were washed 2 times with cold 75% ethanol, airdried and dissolved in 15 μL NaHCO_3_ (0.1M in H_2_O, pH8.3).

#### 3. Coupling NHS-ester dyes to the oligos

The dye labeling reactions included 15 μL Amino-modified oligos in NaHCO_3_ from above and 0.75 μL of succinimide-dye conjugate (20 mM, dissolved in DMSO). The mixture was incubated at room temperature for 2 h in the dark. Oligos were ethanol precipitated by adding to each 15.75 μL reaction mixture: 34.25 μL nuclease free water, 5 μL sodium acetate (3M) and 137.5 μL 100% ethanol; cooling to −80c for 1 h; and centrifuging at 14,000 rpm for 15 min. The now colored pellets were washed 3 times with prechilled 75% ethanol, airdried, and dissolved in 50 μL nuclease free water or Tris-EDTA buffer (10 mM Tris-HCl, 1mM EDTA).

#### 4. Column Purification

To eliminate free fluorescent dyes, the labeled oligos went through column purification. We added 5 μL sodium acetate (3M, pH5.2) and 137.5 μL 100% ethanol to 50 μL dissolved dye-labeled oligos and cooled the mixture at −80C for at least 1 h. The cold mixture was quickly transferred to a PCR-clean up column (Promega, A9282) and centrifuged for 30 s at 14,000g. This step should be done very quickly, as warming of the tube may enhance the solubility of the oligos and reduce their binding to column membrane. Columns carrying the membrane retained probes were washed 2 times with prechilled 80% ethanol and then spun once to remove extra ethanol. 50 μL Nuclease free water or Tris-HCl buffer was added to elute the labeled probes. The probe concentrations were measured by Nanodrop, and used for FISH experiments.

#### Quantification of Degree of Labeling

Degree of Labeling (DOL) of the labeled oligonucleotides was analyzed by UV-Vis spectroscopy using a Nanodrop spectrophotometer and by PAGE.

The DOL of labeled oligos by spectroscopy was estimated using the following formula:

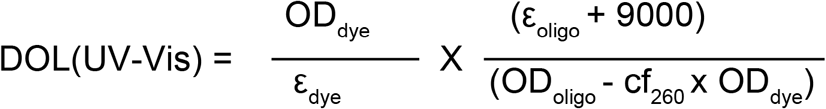

Absorbance at 260 nm (OD_oligo_) and at the dye maximal absorbance (OD_dye_) were measured by Nanodrop and the above formula was used to estimate the properties of the labeled oligonucleotides. ε_dye_ is the extinction coefficient of dye and the cf_260_ represents the relative absorbance of dye at 260 nm. A ε_oligo_ is calculated by taking average of the extinction coefficients of the oligos in the mixture and adding 9000 mol/cm corresponding to the incorporated ddUTP. The DOL of labeled oligos by PAGE was determined using the following formula:

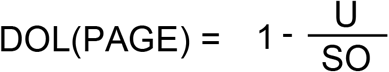

U stands for the band intensity by SYBR staining of the remaining unlabeled oligos after dye labeling, SO stands for the band intensity by SYBR staining of the starting oligos before labeling.

#### Experimental protocol for FISH

C2C12 cells or mouse ES cells were cultured on gelatin coated coverslips. The cells were washed once with ice-cold PBSM (1×PBS, 5mM MgCl_2_), followed by fixation with 4% paraformaldehyde in PBSM for 10 min at room temperature (RT). After a 5-minute wash with ice-cold PBSM, the cells were immersed in 70% ethanol for permeabilization overnight. The ethanol was then removed and 10% formamide in 2 × SSC buffer (300 mM sodium chloride, 30 mM sodium citrate) was added for 2 hours to rehydrate and pre-denature the RNAs. The probes were suspended in hybridization buffer (Biosearch: SMF-HB1-10, 10% formamide added freshly) to a final concentration of 0.5 ng/μL. The coverslips were removed from the dish, dipped on a Kim wipe to remove extra buffer, and then flipped onto droplets of hybridization buffer on parafilm. Hybridization was carried out in humidified chambers at 37C for at least 16 hours. The samples were then washed twice with wash buffer A (Biosearch: SMF-WA1-60) for 30 min at 37C. DAPI (Invitrogen: D1306) was added in the second wash to a final concentration of 0.5 ug/ml. The cells were then washed once with wash buffer B (Biosearch: SMF-WB1-20) for 5 min. The coverslips were mounted in ProLong Gold antifade media (Invitrogen: P36930) overnight before microscopy.

## Acknowledgments

We thank Imre Gaspar, Anne Ephrussi, Tyler Matheny, Roy Parker, Yolanda Markaki, Kathrin Plath, Tamer Sallam and Fangyuan Ding for advice and materials for oligo labeling and FISH; Mohammad Nazim for his critical reading of the manuscript and helpful discussion. This work was supported by NIH Grants R35 GM136426 and R01 GM114463 to DLB, a grant from the W.M. Keck Foundation to DLB, and grants from the Broad Stem Cell Research Center and COVID-19 Research Grant Program at UCLA to DLB.

---

**Table S1.** Reagents and materials required for FISH probe labeling with approximate cost. See detailed information in methods section.

| Reagents | Amount | Price (\$) | Total Reactions | Cost/Reaction (\$) | Cost/ hybridization (\$) |
| --- | --- | --- | --- | --- | --- |
| Oligos (48x) | 200 uM each in ~100ul H2O | 120 | >250 | <0.48 | ~0.015 |
| NHS-ester-Dye | 1 mg | 130-240 | 75 | 1.73-3.2 |  |
| Amino-11-ddUTP | 1 umol | 549 | 100 | 5.49 |  |
| TdT enzyme | 2500 U | 280 | 42 | 6.67 |  |
| Total |  | 1079-1189 |  | 14.37-15.84 |  |

**Table S2.** Chemical structures, catolog numbers and vendors of dyes used in this study.

| Dye-NHS-ester | Structure | Cat.No. |
| --- | --- | --- |
| Quasar570 |  | Biosearch Technology<br>FC-1063S-5 |
| Quasar670 |  | Biosearch Technology<br>FC-1065S-5 |
| ATTO488 |  | Sigma-Aldrich<br>41698-1MG-F |
| ATTO565 |  | Sigma-Aldrich<br>72464-1MG-F |
| ATTO647N |  | Sigma-Aldrich<br>18373-1MG-F |
| Alexa Fluor 568 |  | Fisher Scientific<br>A20003 |
| Alexa Fluor 647 |  | Fisher Scientific<br>A20006 |

